# Effects of genomic location on ectopic integration and gene expression of a reporter gene cassette in *Sulfolobus acidocaldarius*

**DOI:** 10.1101/2025.03.30.646170

**Authors:** Yifei Xu, Andries Ivo Peeters, Indra Bervoets, Marjan De Mey, Rani Baes, Eveline Peeters

## Abstract

In eukaryotes and bacteria, it is well-established that the genomic location of ectopic gene integration influences the expression level due to replication-associated gene dosage effects as well as effects mediated by chromatin organization. In contrast, in archaea, the impact of genomic location on gene expression remained unexplored. Here, we investigated this impact in the model archaeon *Sulfolobus acidocaldarius*, a crenarchaeal species that has a chromatin architecture with mixed eukaryotic-like and bacterial-like features. We aimed to integrate a standardized β-galactosidase (*lacS*) reporter cassette into diverse loci in the genome of *S. acidocaldarius* SK-1 for a comparative analysis. Nine integration mutant strains were successfully obtained, for which qRT-PCR analysis and *lacS* reporter gene assays revealed significant variation in transcriptional and translational expression of the reporter, respectively, demonstrating that genomic location strongly influences gene expression in *S. acidocaldarius*. However, variability in transcription levels and its regulation was shown to be primarily driven by transcriptional activity of neighboring genes, due to the high coding density in the *S. acidocaldarius* genome as well as a lack of insulator elements. Interestingly, translational activity exhibited a more apparent correlation with the distance to the closest origin of replication (*R*^2^ = 0.432) as compared to transcriptional activity (*R*^2^ = 0.026). In conclusion, this study not only provides insights into genome context effects, but also provides inspiration for the future design of genomic knock-in constructions in *S. acidocaldarius*.

## 1 Introduction

Within the Crenarchaeota, species belonging to the Sulfolobales are considered important models for fundamental studies of molecular biological processes in archaea (Lee et al. 2022), also providing evolutionary insights. In addition, they show promise as hosts for biotechnological applications due to their thermoacidophilic lifestyle — growing optimally at a high temperature between 75 and 80 °C and a low pH between 2 and 3 — and their metabolic characteristics, for example chemoorganotrophic metabolism (Crosby et al. 2019). Because of its genetic stability, *Sulfolobus acidocaldarius* is regarded as a chassis species for genetic studies, as well as for genetic engineering for biotechnological purposes (Chen et al. 2005, Wagner et al. 2012, Crosby et al. 2019).

Over the past two decades, a suite of genetic tools has been developed for *S. acidocaldarius* (Albers and Driessen 2008, Wagner et al. 2012, Lee et al. 2022). These tools include electroporation procedures to introduce exogenous DNA, shuttle plasmid vectors derived from native *Saccharolobus* viruses and plasmids, as well as genome engineering approaches based on homologous recombination. Indeed, the native homologous recombination machinery of *S. acidocaldarius* was shown to be sufficiently efficient to enable recombineering (Kurosawa and Grogan 2005, Wagner et al. 2009, Suzuki and Kurosawa 2017). Pyrimidine auxotrophy is commonly used as a selection strategy, with the *pyrEF* operon serving as a selection marker. Selection is performed with uracil and counterselection with 5-fluoroorotic acid (5-FOA), thereby enabling deletion of genomic segments or integration of exogenous DNA with a “pop-in pop-out” approach. This results in the removal of the selection marker, allowing the same marker to be used for multiple subsequent alterations (Wagner et al. 2012).

Using the “pop-in pop-out” method, a repertoire of markerless gene deletion mutants of *S. acidocaldarius* has been constructed, facilitating the study of gene functions (Wagner et al. 2012, Lee et al. 2022). In contrast, the use of this approach for the ectopic integration of a heterologous gene cassette, generating a “knock-in” mutant of *S. acidocaldarius*, has been more scarcely reported (Lee et al. 2022). Nevertheless, in the light of developing *S. acidocaldarius* as a biotechnological host, it is highly relevant to introduce novel genetic traits, such as heterologous enzyme expression, directly into the genome rather than relying on plasmid-based transformations, which might suffer stability issues. In the few reported cases of ectopic gene cassette integration into *S. acidocaldarius*, the genomic location for integration was primarily selected with the aim of minimizing impact on the functioning of genes overlapping or adjacent to the integration site (Zeldes et al. 2019, Lee et al. 2022). For example, the *upsE* locus in *S. acidocaldarius*, encoding UV-inducible pili, was chosen as a target site for ectopic integration of the glucose transporter operon from *Saccharolobus solfataricus*, as *upsE* was previously successfully deleted (Wagner et al. 2012). In another example, a cassette encoding a sulfur oxidation operon was integrated into the *Saci_1149* locus, concomitantly deleting a gene encoding an enzyme in the 3-hydroxypropionic acid/4-hydroxybutyrate (3HP-4HB) pathway, which was shown to be inactive in *S. acidocaldarius* (Zeldes et al. 2019). Finally, a gene cassette encoding a β-xylosidase from *Sa. solfataricus* was integrated into the *pyrEF* locus of *S. acidocaldarius*, which was already disrupted in the host strain (Lee et al. 2022).

In the above case studies, it has not been considered whether or how the genomic location of integration affects gene expression of the integrated gene or operon due to genetic context effects. To our knowledge, this is an outstanding question in Sulfolobales and in archaea in general. In contrast, in eukaryotes it is well-established that overall transcriptional activity varies between chromosomal regions, thereby correlating with gene density and with chromatin organization into discrete compartments (Lieberman-Aiden et al. 2009, Bonev and Cavalli 2016, Rowley and Corces 2018). In bacteria, despite their simpler prokaryotic genome structure, gene expression is also subject to position-dependent effects (Cooke et al. 2019). This has been shown in *Escherichia coli* in various studies (Beckwith et al. 1966; Sousa et al. 1997; Block et al. 2012; Bryant et al. 2014; Brambilla and Sclavi, 2015; Scholz et al. 2022), such as by transposing a β-galactosidase (*lacZ*) reporter gene cassette to different chromosomal locations (Sousa et al. 1997). It was found that gene expression increases with an increasing proximity to the origin of replication, an effect linked to replication-associated gene dosage (Beckwith et al. 1966). Additional local effects, such as differences in genome accessibility to RNA polymerase, chromatin structure and organization, nucleoid-associated proteins (NAPs) and DNA topology were also observed (Block et al. 2012; Bryant et al. 2014; Brambilla and Sclavi, 2015; Scholz et al. 2022). For instance, also in *E. coli*, a GFP reporter cassette exhibited gene expression differences of up to 300-fold depending on chromosomal location (Bryant et al. 2014).

Intriguingly, while archaea share a similar gene organization to bacteria, typified by operonic structures and high coding density, their chromatin is structured using a combination of bacterial and eukaryotic mechanisms. In *S. acidocaldarius*, the chromosome is three-dimensionally organized into a higher-order structure reminiscent of eukaryotic chromatin, forming two chromosomal compartments: compartment A is characterized by a higher transcriptional activity, containing essential genes and replication origins and compartment B is characterized by a lower transcriptional activity and is enriched in non-essential genes and transposons (Takemata et al. 2019, Pilatowski-Herzing et al. 2024). Unlike most archaea, Crenarchaea, including *S. acidocaldarius*, lack condensin, a protein involved in chromatin organization (Pilatowski-Herzing et al. 2024). Instead, Sulfolobales harbor an SMC protein named coalescin (ClsN), which is enriched in the B compartment and associated with gene silencing (Takemata et al. 2019). On a smaller scale, the chromatin of *S. acidocaldarius* and other Sulfolobales species is organized by a diverse repertoire of small (7–12 kDa) NAPs, including Cren7, Sul7, Alba, Sul10a and Sul12a, which contribute to chromatin structuring through diverse DNA interactions, including DNA kinking, compaction, bridging, looping and supercoiling (Grote et al. 1986; Lurz et al. 1986; Choli et al. 1988; Baumann et al. 1994; Lopez-Garcia et al. 1998; Bell et al. 2002; Guo et al. 2008; Driessen et al. 2016; Kalichuk et al. 2016; Zhang et al. 2020; Lemmens et al. 2022).

In this study, we address the hypothesis that the genomic location of ectopic integration affects gene expression in *S. acidocaldarius*, similarly as in eukaryotes and bacteria. Specifically, we aim to explore how a location within each of the higher-order A or B chromatin compartments in *S. acidocaldarius* affects gene expression, given that they show significant differences in global transcriptional activity in the native transcriptome (Takemata et al. 2019). This will be achieved by integrating a constitutive β-galactosidase (*lacS*) gene reporter cassette into different genomic locations which are selected based on differences in chromatin compartment and proximity to a replication origin, as well as the expression levels and predicted essentiality of neighboring genes.

## 2 Materials and methods

### 2.1 Microbial strains and cultivation conditions

The uracil-auxotrophic strain *S. acidocaldarius* SK-1 (Suzuki and Kurosawa 2016) and all derived mutant strains were cultivated in liquid in basic Brock medium (Brock et al. 1972) supplemented with 0.1% (w/v) N-Z-Amine, 0.2 % (w/v) sucrose and 20 µg ml^-1^ uracil, acidified to pH 3.0 with sulfuric acid. Liquid *S. acidocaldarius* cultures were incubated at 75 °C while shaking. Growth was monitored by measuring optical density at 600 nm (OD^600^). For growth on solid medium, Brock medium was supplemented with 1.5 mM CaCl_2_, 5 mM MgCl_2_ and 0.6% Gelrite as a solidifying agent (Wagner et al. 2012). For selection of marker-containing integration mutants (“pop-in” step), the medium was solely supplemented with 0.1% (w/v) N-Z-Amine, 0.2% (w/v) sucrose, while for selection of markerless knock-in mutants (“pop-out” step), the medium was supplemented with 0.1% (w/v) N-Z-Amine, 0.2% (w/v) sucrose, 20 µg ml^-1^ uracil and 200 µg ml^-1^ 5-FOA. Solid medium was prewarmed to 75 °C before performing plating (Wagner et al. 2012). Nutrient starvation was performed based on (Haurat et al. 2017). Cells were cultivated until reaching OD_600_ 0.4 and collected by centrifugation at 4000 *g* for 10 minutes at room temperature. The cell pellet was resuspended in fresh prewarmed Brock medium supplemented with 20 µg ml^-1^ uracil lacking N-Z-Amine and sucrose and incubated at 75 °C for 4 hours while shaking.

*E. coli* strain DH5α was used for cloning and for propagation of plasmid constructs and was cultivated in Lysogeny Broth (LB) medium supplemented with 50 µg ml^-1^ ampicillin, if required. For selection of transformants from Golden Gate cloning, LB medium was used in which NaCl was omitted and 5% (w/v) sucrose was added. All strains used in this work are provided in **Supplementary Table S1**.

### 2.2 DNA manipulations and molecular cloning

Gibson assembly (Gibson et al. 2009) was used to construct a Golden Gate destination vector named pYX2304. To this end, the backbone of suicide plasmid vector pSVA431 (Wagner et al. 2012) was amplified using oligonucleotides YX123 and YX124 (**Supplementary Table S2**) and the *sacB* gene was amplified from the pRN1-carrier_2_paqci plasmid vector using primers YX125 and YX126. Polymerase Chain Reaction (PCR) amplification was performed with KAPA HiFi DNA polymerase (Roche) using 10 ng plasmid DNA and 20 pmol of each primer in a 50 µL reaction with following PCR conditions: 3 minutes at 95 °C, 35 cycles of 20 seconds at 98 °C, 15 seconds at 60 °C and 1 minute/kb at 72 °C, followed by a final step of 5 minutes at 72 °C. PCR amplification was verified by agarose gel electrophoresis and products were purified using the Wizard® SV Gel and PCR Clean-Up System (Promega). Next, Gibson assembly was performed by incubating 100 ng of amplified pSVA431 backbone with an equimolar amount of the *sacB* gene fragment in NEBuilder HiFi DNA Assembly Master Mix (New England Biolabs) at 50 °C for 1 hour, followed by heat-shock transformation into chemically competent *E. coli* DH5α. Transformants were screened by colony PCR using primers YX127 and YX128 and the sequence of the obtained construct was verified by Sanger sequencing (Eurofins Genomics).

The pYX2304 destination vector was subsequently used to construct all suicide knock-in plasmid vectors with Golden Gate assembly (Engler et al. 2008). For each of the targeted genomic locations, primers (**Supplementary Table S2**) were designed to enable amplification of fragments of approximately 650 bp up- and downstream of the target site, respectively. These PCR reactions were performed using KAPA HiFi DNA polymerase (Roche) with 10 ng genomic DNA (gDNA) of *S. acidocaldarius* SK-1, which was extracted from liquid culture employing a QuickPick SML gDNA kit with magnetic bead purification (BioNobile). The *lacS* reporter cassette was PCR-amplified using KAPA HiFi DNA polymerase (Roche) with primers YX133 and YX134 and pYX2301-UPDD plasmid DNA as a template. The latter was derived from pSVA431 by replacing the P*malE* promoter driving *lacS* expression with a synthetic constitutive promoter named P_Ec10_Sa1_. This was realized via Gibson assembly, with the 56-bp P_Ec10_Sa1_ promoter sequence being introduced by primer YX091 (**Supplementary Table S2**). Golden Gate assembly was performed in 20-µl reactions consisting of 2 µl T4 ligase buffer (ThermoFisher), 1 µl T4 ligase (ThermoFisher), 1 µl PaqCI (Bioké), 0.3 µl PaqCI activator (Bioké), 100 ng pYX2304 plasmid DNA and equimolar amounts of the fragments. Reaction mixtures were subjected to following conditions: 50 cycles of 3 minutes at 37 °C, 4 minutes at 16 °C, followed by a final step of 5 minutes at 37 °C and 5 minutes at 60 °C. Subsequently, reaction mixes were heat-shock transformed into chemically competent *E. coli* DH5α and transformants were screened by colony PCR using customized primers (**Supplementary Table S2**). Finally, the sequence of the obtained construct was verified by Sanger sequencing (Eurofins Genomics). An overview of all plasmids constructed and used in this work is provided in **Supplementary Table S3**.

### 2.3 Construction of *S. acidocaldarius* knock-in mutant strains

Construction of *S. acidocaldarius* markerless knock-in mutant strains was performed using a classical “pop-in pop-out” strategy as described (Wagner et al. 2012, Suzuki and Kurosawa 2016). Competent *S. acidocaldarius* SK-1 cells were prepared by harvesting cells from a 100-ml culture at an OD_600_ of 0.2 through centrifugation at 6574 *g* for 20 minutes, followed by two washing steps in 20 mM sucrose and a final resuspension of the cells in this buffer, reaching a final concentration of 2×10^10^ cells ml^-1^. Genetic transformation was performed with a Gene Pulser II electroporator (Bio-Rad) at 1.5 kV, 25 mF and 600 W with Gene Pulser 1-mm cuvettes (Bio-Rad) (Wagner et al. 2012). After 5 days of incubation, transformant colonies were screened for β-galactosidase activity by spraying with 5 mg/ml 5-bromo- 4-chloro-3-indolyl-beta-D-galacto-pyranoside (X-Gal) (ThermoFisher), followed by an incubation at 75 °C for 30 minutes. Positive “blue” integrants were further confirmed by PCR analysis and were cultivated in liquid culture without addition of uracil. Fifty µL of liquid culture were subsequently plated on Gelrite plates with uracil (20 μg/ml) and 5-FOA (200 μg/ml) for the second selection of “pop-out” of the vector backbone by homologous recombination and incubated for 5 days at 75 °C. Transformant colonies were screened for β-galactosidase activity as described previously and successful integrants were confirmed through PCR analysis.

### 2.4 Quantitative reverse transcriptase PCR

*S. acidocaldarius* cells were cultivated in a liquid medium until reaching an OD_600_ of between 0.4 and 0.5 (exponential growth phase) or OD_600_ of 1.0 (stationary growth phase). Culture samples of 4 mL were centrifuged for 15 minutes at 4000 *g* and pellets were stabilized using an equal volume of RNAprotect (Qiagen) and subjected to RNA extraction using an SV Total RNA Isolation System kit (Promega), followed by removal of residual genomic DNA using a Turbo DNase kit (Ambion Life Technologies). Next, cDNA was prepared from 1 mg RNA using a Go-Script Reverse Transcriptase kit (Promega). Three biological replicates were included in this experiment, except for the stress condition, which was conducted without replication.

Quantitative reverse transcriptase PCR (qRT-PCR) was performed to determine relative transcriptional gene expression levels of *lacS* employing *tbp* (*saci_1336*) as a reference gene. qRT-PCR primers were designed with Primer3 software (Untergasser et al. 2012) (**Supplementary Table S2**) and tested for efficiency using *S. acidocaldarius* gDNA as a template. Twenty-µL qRT-PCR reactions were performed in an iCycler qPCR device (Bio-Rad) with each reaction mixture containing 1 ml 10-fold diluted cDNA, 1.6 pmol of each primer and GoTaq qPCR master mix (Promega). The following PCR conditions were used: 3 minutes at 95 °C and 40 cycles of 10 seconds at 95 °C and 30 seconds at 55 °C. Quantification cycle (C_T_) values were determined with iQ5 software (Bio-Rad). For each gene and biological replicate, three technical replicates were performed, for which the average C_T_ value was determined. Relative expression levels were normalized with respect to the reference gene and ratios were calculated using the ΔΔC_T_ method relative to the strain displaying the lowest expression level. Graphpad Prism 9 was employed to perform a two tailed, one sample T test for statistical analysis.

### 2.5 Western blotting

Four ml of *S. acidocaldarius* cells, cultivated until an OD_600_ of 0.4, were pelleted by centrifugation at 4000 *g* for 15 minutes at 4 °C. The pellet was resuspended in 200 mL extraction buffer (50 mM Tris-HCl, 50 mM NaCl, 15 mM MgCl_2_, 1 mM DTT) to which 0.1% Triton X-100 was added. Cells were incubated at 4 °C for 30 minutes while rotating for lysis. Total protein concentration was determined using Bradford Assay solution (TCI Chemicals), using BSA for the generation of the standard curve.

Normalized protein extracts (30 μg) were mixed with LDS (ThermoFisher) and subjected to sodium dodecyl sulfate poly-acrylamide gel electrophoresis (SDS-PAGE) using 4-12% NuPAGE™ Bis-Tris Mini Protein gels (ThermoFisher) and NuPAGE™ MES SDS Running Buffer (ThermoFisher). After SDS-PAGE, proteins were electroblotted onto a polyvinylidene difluoride (PVDF) membrane (Bio-Rad) using a Trans-Blot Turbo transfer system (Bio-Rad) operated at 1.3 A, up to 25 V for 7 minutes. The membrane was then incubated in Phosphate Buffer Saline (PBS) buffer with 5% (v/v) milk and 0.1 % (v/v) Tween20 for 1 hour at room temperature followed by an overnight incubation at 4 °C with anti-His-tag mouse antibody (Proteintech, Cat No. 66005-1-Ig) diluted at 1:5000 in the same buffer. Excess primary antibody was washed away with PBS buffer containing 0.1% Tween20, followed by incubation with a secondary goat anti-mouse HRP-conjugated antibody (Proteintech, Cat No. SA00001-1) diluted at 1:5000 in the same buffer. After 1 hour incubation at room temperature, unbound antibodies were washed away with PBS buffer containing 0.1% Tween20, twice,and PBS buffer for final wash. Visualization was performed with a Pierce™ ECL Western Blotting Substrate kit (ThermoFisher) and an ImageQuant800 imager (Cytiva).

### 2.6 β-galactosidase reporter gene assays

β-galactosidase assays were performed as described (van der Kolk et al. 2020) with some modifications. The σ-nitrophenyl-β-D-galactopyranoside (ONPG) conversion rate was measured spectrophotometrically at 410 nm in a microplate reader (Infinite M Nano+, TECAN) every 5 minutes, while being incubated at 42 ºC for 4 hours. The slope of the conversion curves was taken as the value for β-galactosidase activity, which is a proxy for LacS expression. For each sample, the assay was performed in triplicate. The slopes were determined by fitting a Monod-type model with smoothing to the conversion data:

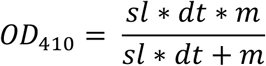

with

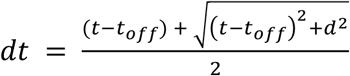

where *sl* = slope, *m* = maximum value, *t* = time, *t*_*off*_ = offset and *d* = period around the offset that the transition takes place. The slope was determined by fitting the model to ranges of datapoints from t_0_- t_max_ to t_0_-t_5_. Local minima of the fit error of these fits were then searched and from those fits, the slope of the fit resulting in the slope with the lowest standard error was selected. When the resulting fit was not deemed adequate, the average value of the fitted *d* parameter of the other technical and/or biological replicates was used as fixed value for a subsequent fitting. The geometric mean of the slopes was calculated, together with the geometric standard error. If the fit error or error from the technical replicates was larger than the geometric standard error, that one was used instead.

## 3 Results

### 3.1 Establishment of an efficient suicide vector cloning system for construction of knock-in mutant strains

In this study, we used a knock-in suicide plasmid vector that is based on the pSVA431 plasmid vector harboring a *Sa. solfataricus pyrEF* gene cassette and a pGEM-T Easy cloning vector containing an ampicillin resistance cassette and f1 origin of replication for *E. coli* (Wagner et al. 2012). To facilitate the construction of knock-in suicide vectors, we have chosen to employ a one-step Golden Gate assembly approach. To this end, pSVA431 was converted into a Golden Gate destination vector named pYX2304 (**Figure 1a**). This vector combines the pSVA431 backbone with a *sacB* gene cassette flanked by type IIS restriction sites. The *sacB* gene encodes levansucrase, an enzyme that converts sucrose into levans, which accumulates in the *E. coli* periplasm causing toxicity (Gay et al. 1985). During cloning, it can thus be used as a negative selection marker in presence of sucrose, enabling the selection of transformants harboring a vector in which the *sacB* cassette is excised and replaced by the fused target up- and downstream regions.

**Figure 1.**
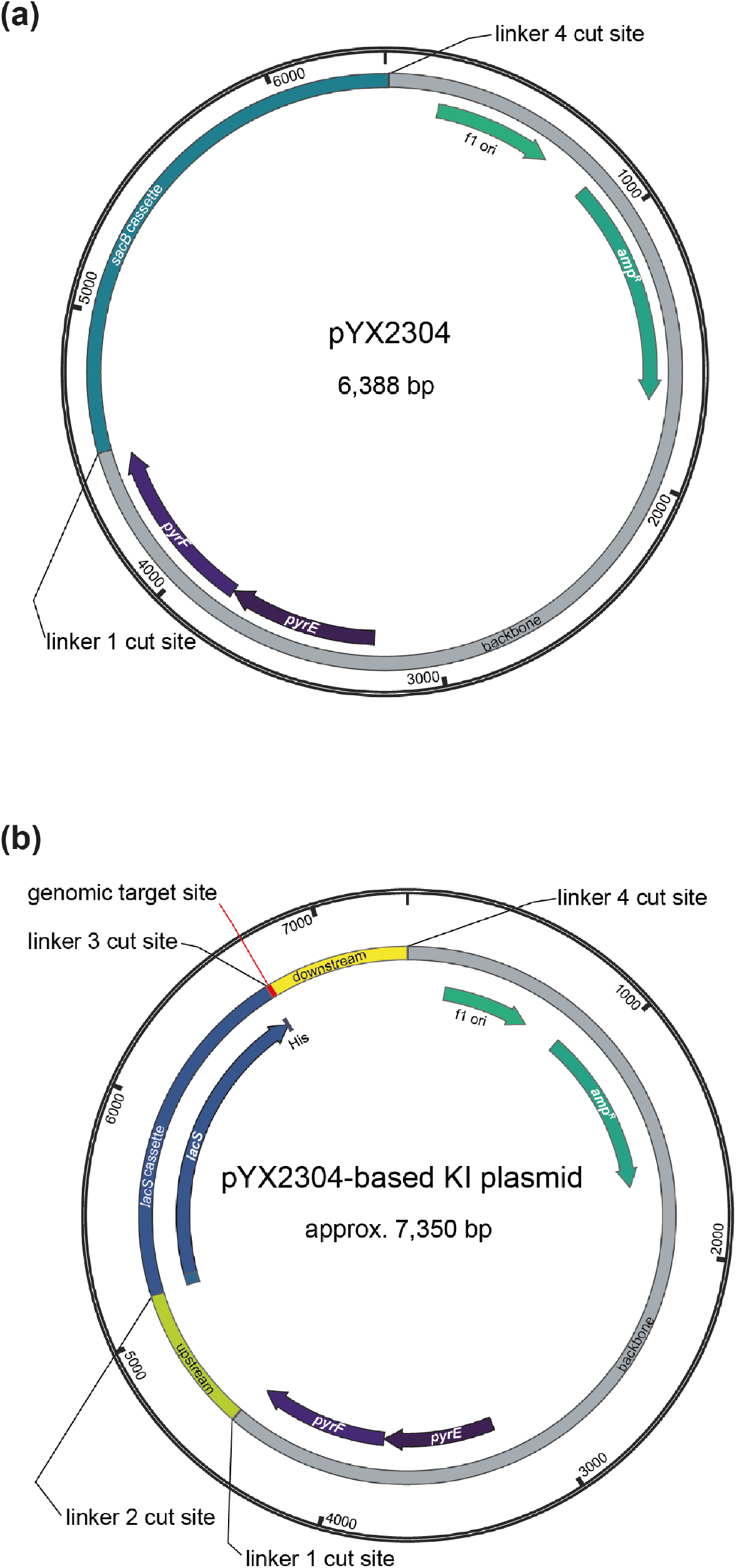
Plasmid maps of the Golden Gate destination vector pYX2304 (**a**) and the derived suicide knock-in (KI) plasmid vector (**b**) with indication of the different type IIS restriction sites with the following cut sites: linker 1 = 5’-AGGA-3’, linker 2 = 5’-TGAT-3’, linker 3 = 5’-GATG-3’ and linker 4 = 5’-GCAG-3’.

The designed knock-in suicide plasmid vector (**Figure 1b**) can be used to create markerless knock-in mutant strains with a single-crossover recombination approach, employing the *pyrEF* cassette as a selection marker (Wagner et al. 2012) and the uracil auxotrophic *S. acidocaldarius* SK-1 (Suzuki and Kurosawa 2016) as a host strain (**Figure 2**). This enables a classical “pop-in pop-out” scheme in which, in a first selection step directly following transformation, cultivation on solid medium lacking uracil results in selection of integrant mutant strains, with the entire plasmid vector being integrated into the genome (“pop-in” step) (Wagner et al. 2012). Successfully obtained integrant mutant strains are then subjected to a second selection step, which entails a counterselection because of the presence of uracil and 5-FOA, resulting in the selection of strains that have undergone a second single-crossover recombination event (“pop-out” step), leading either to the wild-type genomic sequence or to the desired knock-in mutant sequence (**Figure 2**).

**Figure 2.**
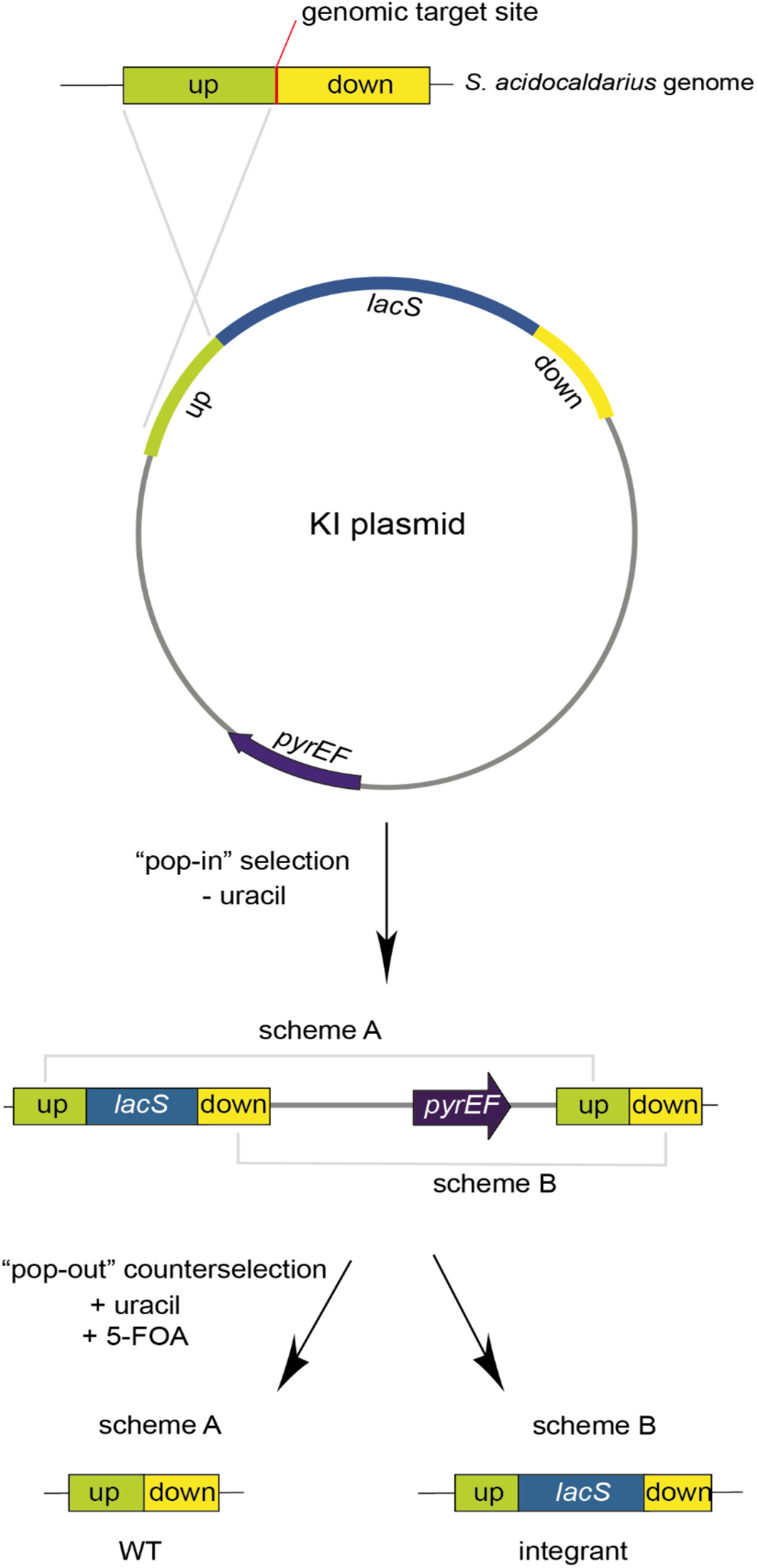
Flowchart of the “pop-in pop-out” approach used for the construction of knock-in mutant strains in *S. acidocaldarius* SK-1. The genomic target site is indicated with a red line.

The *lacS* reporter cassette consisted of a *lacS* gene from *Sa. solfataricus* fused to a C-terminal 6xHis-tag and under control of a synthetic *Sulfolobus* promoter named P_Ec10_Sa1_, which is based on the native *Saci_2137* promoter, driving expression of a putative aminotransferase (Liu et al., 2014).

### 3.2 Efficiencies in genome integration and in obtaining markerless knock-in mutant strains

We selected eleven target positions for insertion within the *S. acidocaldarius* SK-1 genome, based on i) variations in the relative distance of the site to the nearest origin of replication, ii) transcriptional expression levels of nearby or overlapping genes under physiological conditions (Baes et al. 2023) and iii) predicted essentiality assessed based on homology with *Saccharolobus islandicus* genes (Zhang et al. 2018) (**Figure 3a** and **Table 1**). Additionally, the target sites were chosen to be evenly distributed between the two chromosomal compartments A and B (**Figure 3a** and **Table 1**). In all cases with one exception (*cmp*), the reporter gene cassette was specifically targeted to intergenic regions to minimize unintended effects on the expression of adjacent genes (**Figure 3b** and **Supplementary Figure S1**). Intergenic regions include those located between genes transcribed in the same direction, whether part of an operon (*e*.*g. ccc1*) or not (*e*.*g. sdhC*), as well as between convergently (*e*.*g. cmp*) or divergently transcribed genes (*e*.*g. acad*). The insertion positions and corresponding knock-in strains were named based on the nearest gene (**Table 1**).

**Table 1.**
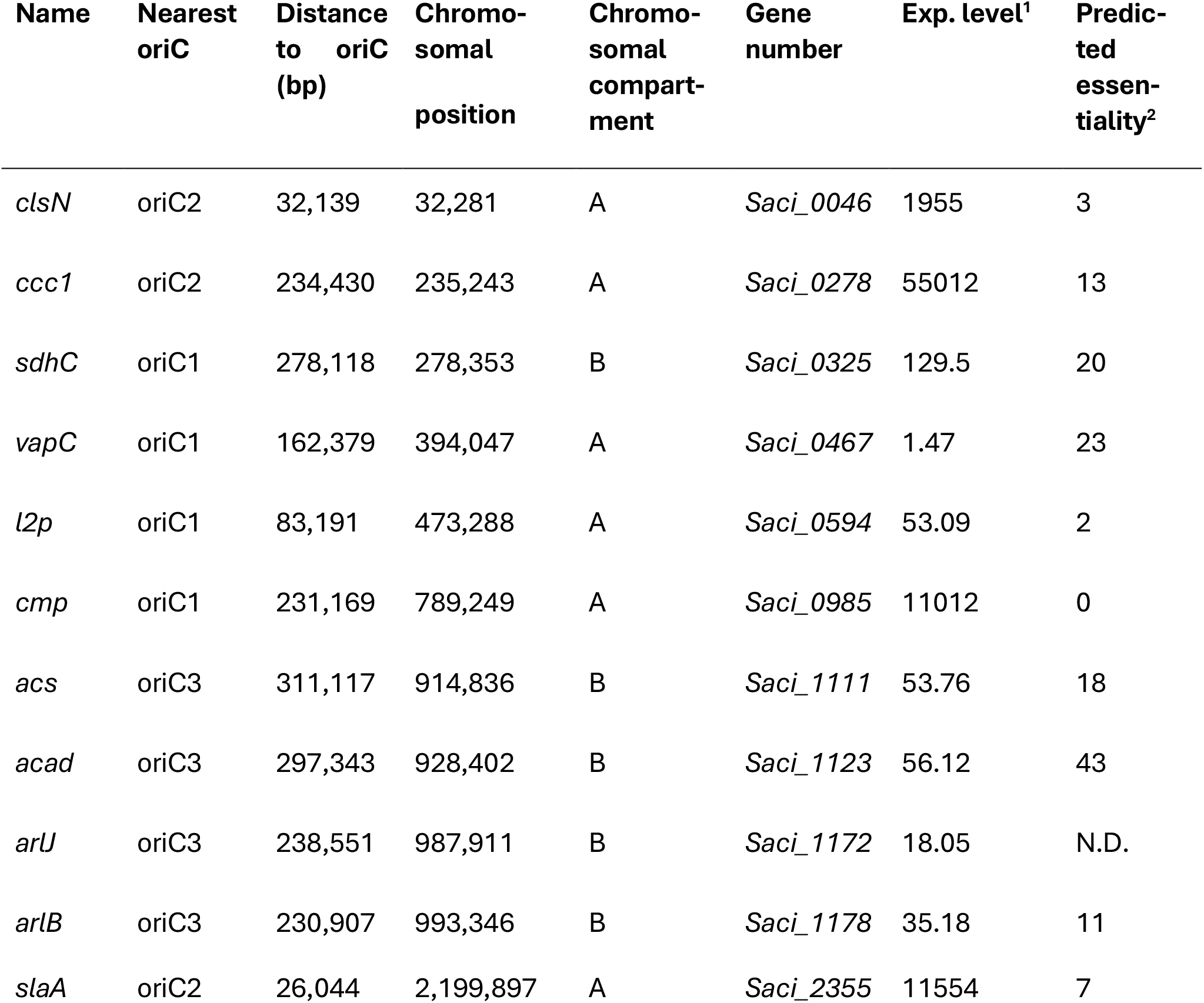
Overview of the eleven insertion positions, with indication of name, nearest origin of replication, distance to nearest origin of replication, chromosomal position, chromosomal compartment, as well as gene number, annotation, expression level (Exp. level) and predicted essentiality of the nearest gene (Zhang et al. 2018). All information in this table is based on alignment with the *S. acidocaldarius* DSM639 genome sequence (Chen et al. 2005). ^1^Expression level is expressed in CPM value (Baes et al. 2023). N.D. indicates not determined. ^2^Predicted essentiality is expressed on a scale between 0 and 48 with the latter the highest essentiality.

**Figure 3.**
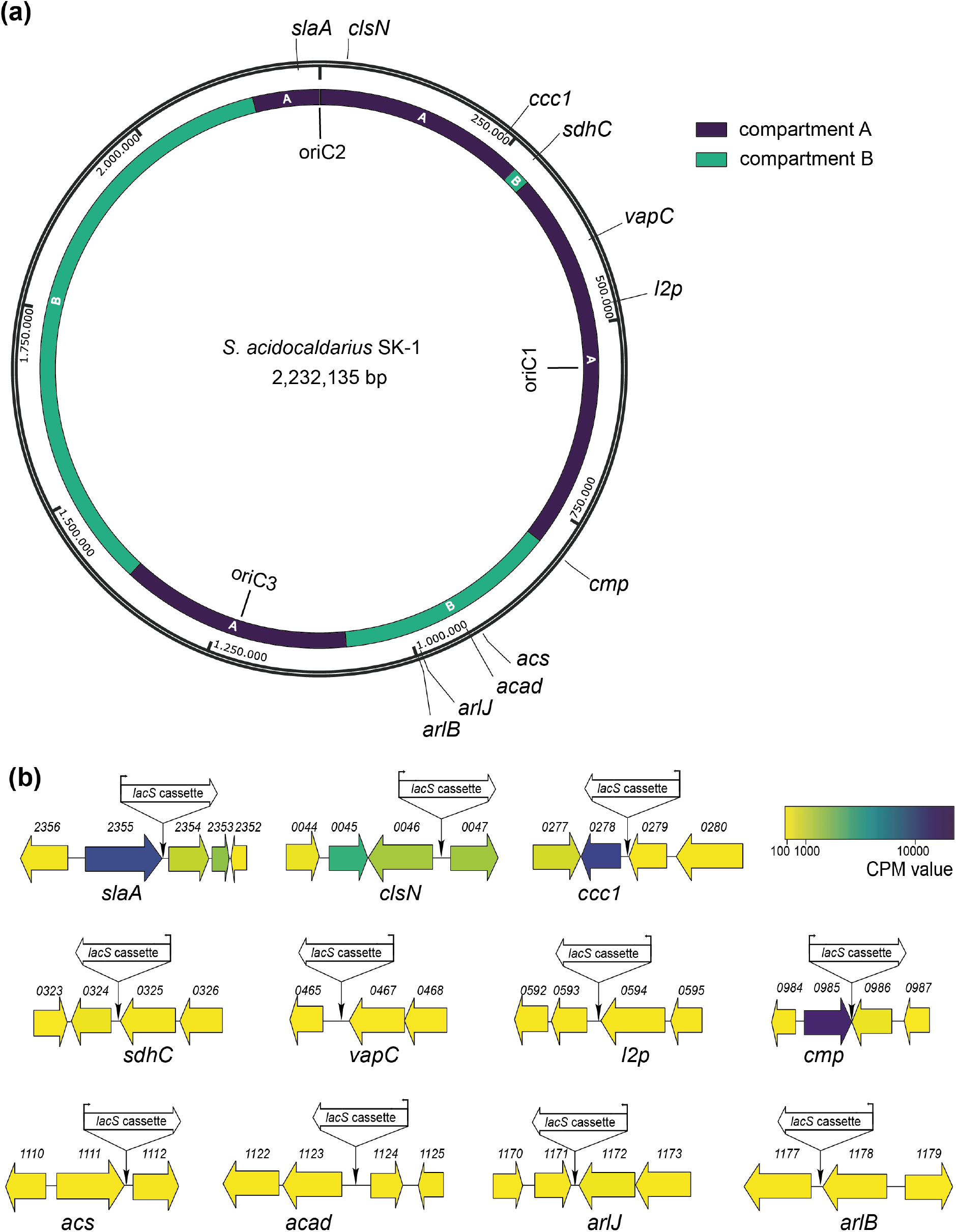
Overview of 11 selected genomic target sites for integration of a *lacS* cassette. (**a**) Map of the *S. acidocaldarius* SK-1 genome sequence with indication of genomic position, replication origins (oriC1, oriC2 and oriC3) (Duggin et al. 2008) and target sites (with naming according to **Table 1**). Chromosome compartments A and B (Takemata et al. 2019) are color-indicated. (**b**) Genetic organization in the genomic environment of the 11 target sites with indication of the target site and the orientation of the *lacS* cassette. Numbers refer to gene locus tags, with xxxx referring to *Saci_xxxx*. Gene arrows are color-coded according to expression levels (CPM value) (Baes et al. 2023).

All genetic constructs were transformed into *S. acidocaldarius* SK-1 followed by a selection in uracil-free medium, yielding high numbers of colony-forming units (CFUs). Upon X-gal treatment, enabling colorimetric detection of *lacS*^+^ colonies, it appeared that only a small fraction of CFUs represented “pop-in” *lacS*^+^ integrants (**Figure 4a**). The nature of the “background” *lacS*^*-*^ CFUs is unknown. On average, the fraction of “pop-in” *lacS*^*+*^ CFUs was higher for locations in compartment A as compared to compartment B (an average of 0.56 % *versus* 0.22 %) (**Figure 4a**). Transformations that yielded a higher “pop-in” *lacS*^+^ integrant fraction, such as for *slaA* and *ccc1* knock-in constructs, were possibly characterized by more a favorable genomic recombination as compared to transformations with knock-in constructs like *acad* or *arlJ*.

**Figure 4.**
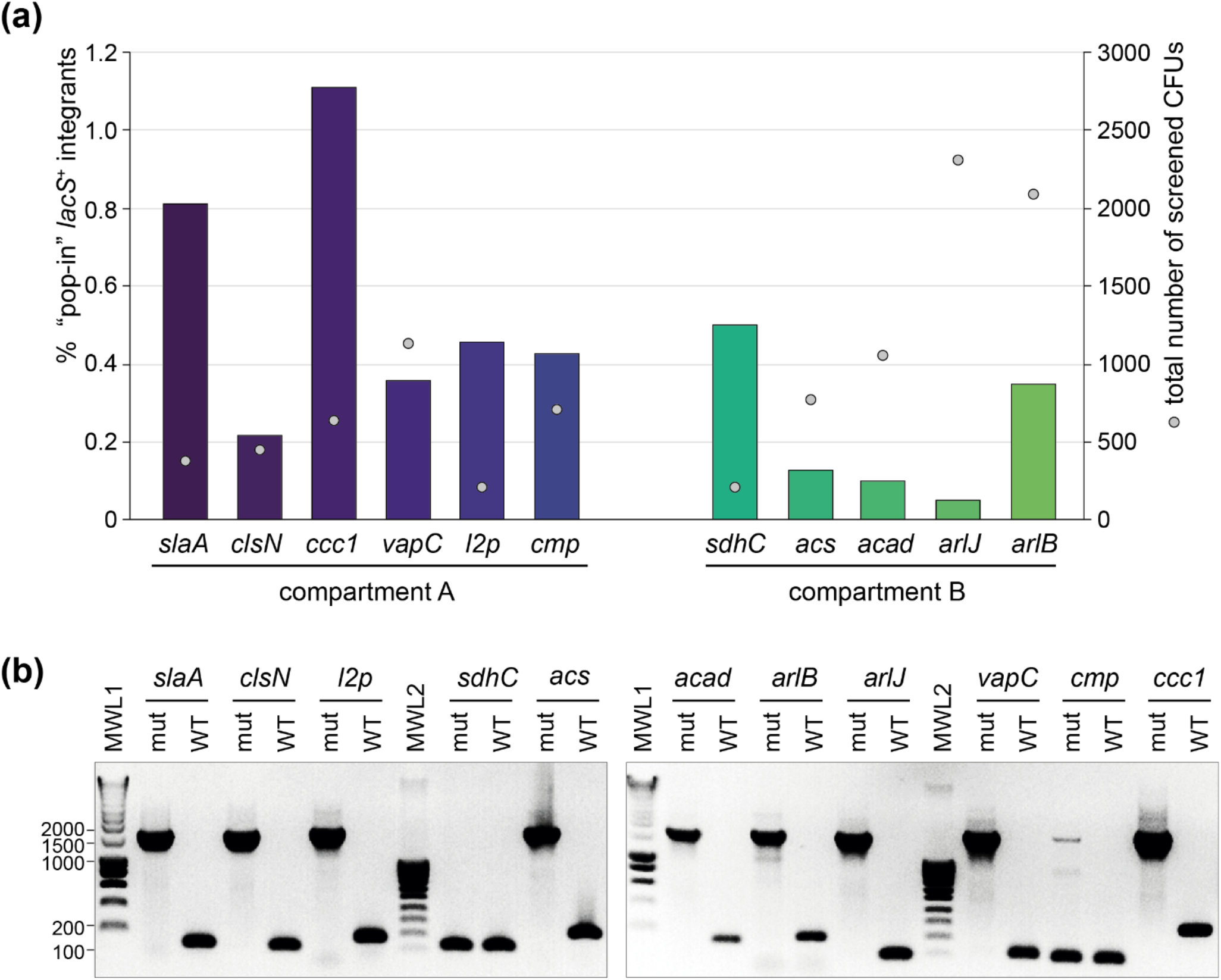
Construction of the *lacS*^*+*^ knock-in mutant strains. (**a**) Percentage of “pop-in” *lacS*^*+*^ integrant CFUs after the “pop-in” construction step, depicted in bars for each of the intended knock-in strains (**Table 1**), grouped according to their chromosome compartment. This percentage is based on X-gal colorimetric screening and expressed with respect to “background” CFUs. Sphere symbols indicate the total number of screened CFUs for each of the knock-in constructions. (**b**) Colony PCR of selected “pop-out” *lacS*^*+*^ integrant strains to verify integration of the *lacS* cassette, leading to an amplicon of approximately 1700 bp. WT = wild type; mut = “pop-out” *lacS*^*+*^ integrant; MWL = molecular weight ladder.

Purified “pop-in” integrant strains were subsequently cultivated on 5-FOA- and uracil-containing plates to select for “pop-out” *lacS*^+^ integrant strains, which still contained the *lacS* cassette, but with the remainder of the vector removed by a crossover event (**Figure 1b**). For all knock-in constructions, with the exception of *sdhC* and *cmp*, “pop-out” *lacS*^+^ integrant strains were easily obtained and the correct integration of the *lacS* cassette was verified by colony PCR (**Figure 4b**). In contrast, for the construction of the *sdhC* and *cmp* knock-in strains, colony PCR revealed partial or full presence of the wildtype genotype and despite multiple selection attempts, we failed in obtaining these knock-in strains. For the *cmp* knock-in strain this could be possibly attributed to the disruption of the open reading frame (ORF) of the adjacent *saci_0956* gene, encoding a putative glycosyltransferase (**Supplementary Figure S1**).

### 3.3 Transcriptional gene expression analysis of *lacS* in the different knock-in strains

A qRT-PCR approach was used to evaluate transcriptional expression levels of the *lacS* reporter gene in the different knock-in strains (**Figure 5**). Although all strains were cultivated in identical conditions and despite *lacS* being under control of the same constitutive promoter, large variations in transcriptional levels were observed (**Figure 5a**). Relative transcription is calculated with respect to the knock-in strain displaying the lowest *lacS* transcription, *arlJ*.

**Figure 5.**
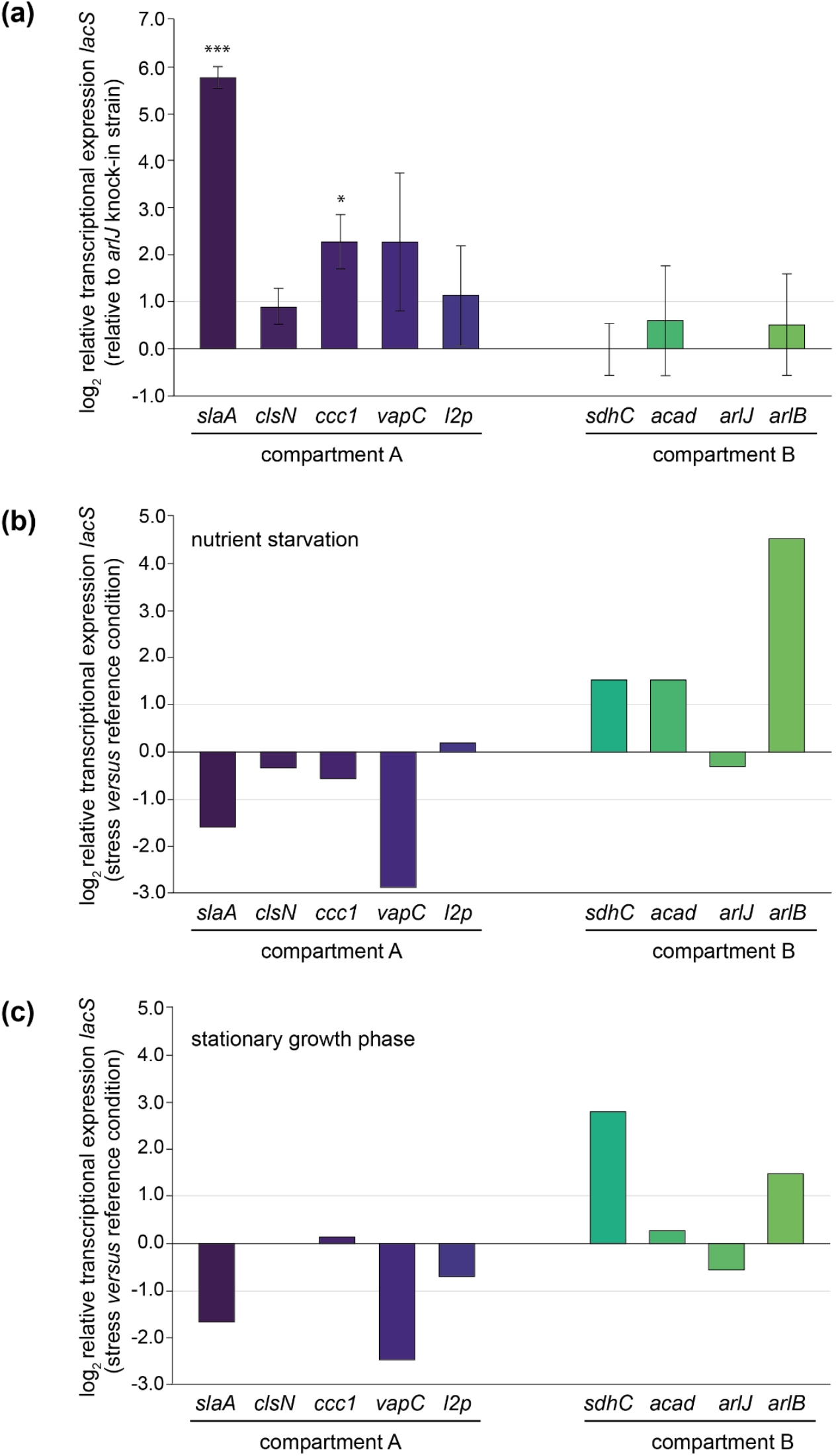
Relative transcriptional expression levels of *lacS* as monitored by qRT-PCR analysis. (**a**) Relative transcriptional *lacS* expression in each of the knock-in strains relative to the *arlJ* knock-in strain, grouped according to the chromosomal compartment. Experiments have been performed in biological triplicates with error bars representing standard deviations. Statistically significant differential expression was determined by a one-sample T-test (* = p < 0.05, ** = p < 0.01, *** = p < 0.001). (**b**) Relative transcriptional *lacS* expression in nutrient starvation conditions *versus* the reference growth condition in each of the knock-in strains. **c**. Relative transcriptional *lacS* expression in stationary *versus* exponential growth phase.

Transcriptional levels were observed to be highest in the *slaA* knock-in strain, 55-fold higher than in the *arlJ* knock-in strain. This could be attributed to the target site position, immediately downstream of the *slaA* stop codon, making it highly probable that *slaA* transcription, which is known to be constitutively expressed at high level (Baes et al. 2023), runs through to the *lacS* cassette. The same applies for other mutant strains that display significantly higher transcription levels with respect to the *arlJ* knock-in strain (**Figure 5a**). For the *vapC* knock-in strain, in which *lacS* is transcribed at a 6.2-fold higher level, the *lacS* cassette was integrated downstream of the *vapC* gene preceding a putative terminator site (**Figure 5a** and **Supplementary Figure S1**). For the *ccc1* knock-in strain, in which *lacS* is transcribed 5.2-fold higher than in the *arlJ* knock-in strain, the target site is located in between two operonic genes (**Figure 5a** and **Supplementary Figure S1**).

Only in the *clsN* and *acad* knock-in strains, the target sites are located in regions where no read-through transcription from adjacent genomically encoded transcription units is expected, namely in an intergenic region in between the two divergently oriented putative promoters (**Supplementary Figure S1**). For these strains, transcriptional expression of *lacS* is expected to be driven solely by the P_Ec10_Sa1_ promoter and indeed, both strains have very similar transcriptional levels with respect to the the *arlJ* knock-in strain, namely 1.88-fold and 1.86-fold for *clsN* and *acad*, respectively (**Figure 5a**).

Although a trend was observed of compartment A-targeted knock-in strains having higher transcriptional expression levels than compartment B-targeted knock-in strains, this cannot be linked to a global effect, but rather to the expression levels of adjacent genes, which are not insulated from the *lacS* cassette. To a certain extent, a correlation was observed between previously detected expression levels of the adjacent gene (Baes et al. 2023) expected to cause read-through transcription and the relative *lacS* transcriptional level (**Supplementary Table S4**). Given that for the two “insulated” knock-in strains *clsN* and *acad*, which both have similar transcriptional expression levels and of which the *clsN* target site is located in compartment A and the *acad* target site in compartment B, a global gene silencing effect was not apparent.

### 3.4 Transcriptional regulation of *lacS* in the different knock-in strains

Transcriptional expression of *lacS* was also monitored in response to stress conditions, either nutritional starvation (**Figure 5b**) or stationary phase growth (**Figure 5c**). Although the P_Ec10_Sa1_ promoter is expected to drive transcription of *lacS* in a constitutive manner, differential transcription was observed in several knock-in strains, either positively or negatively. Trends are similar for both nutritional starvation and stationary growth phase, suggesting that the stress signals experienced by the cells in these conditions is similar. In the *slaA* and *vapC* knock-in strains, a transcriptional downregulation of *lacS* is observed, while in the *sdhC* and *arlB* knock-in strain, an upregulation is observed (**Figures 5b** and **5c**). For the latter strain, the target site of the cassette is directly downstream of the *arlB* ORF (**Supplementary Figure S1**) and assumed to be subjected to the same transcriptional regulatory mechanisms.

In agreement to the stable constitutive relative transcription levels, little or no transcriptional regulation was observed for the “insulated” knock-in strains *clsN* and *acad* (**Figures 5b** and **5c**).

### 3.5 Translational level and activity analysis of LacS in the different knock-in strains

To assess the abundance and activity of LacS-6xHis on the protein level, we performed anti-6xHis western blotting, as well as ONPG-based LacS activity assays (**Figure 6**). Both approaches demonstrated a variability among the knock-in strains with compartment A strains showing a relatively higher average activity as compared to compartment B strains (an average of 2.412*x*10^−4^ versus 1.419*x*10^−4^) (**Figure 6b**), with the *clsN* and *slaA* knock-in strains seemingly exhibiting the highest activity, together with the *vapC* knock-in strain. These results suggest a higher translational efficiency for genes positioned in compartment A.

**Figure 6.**
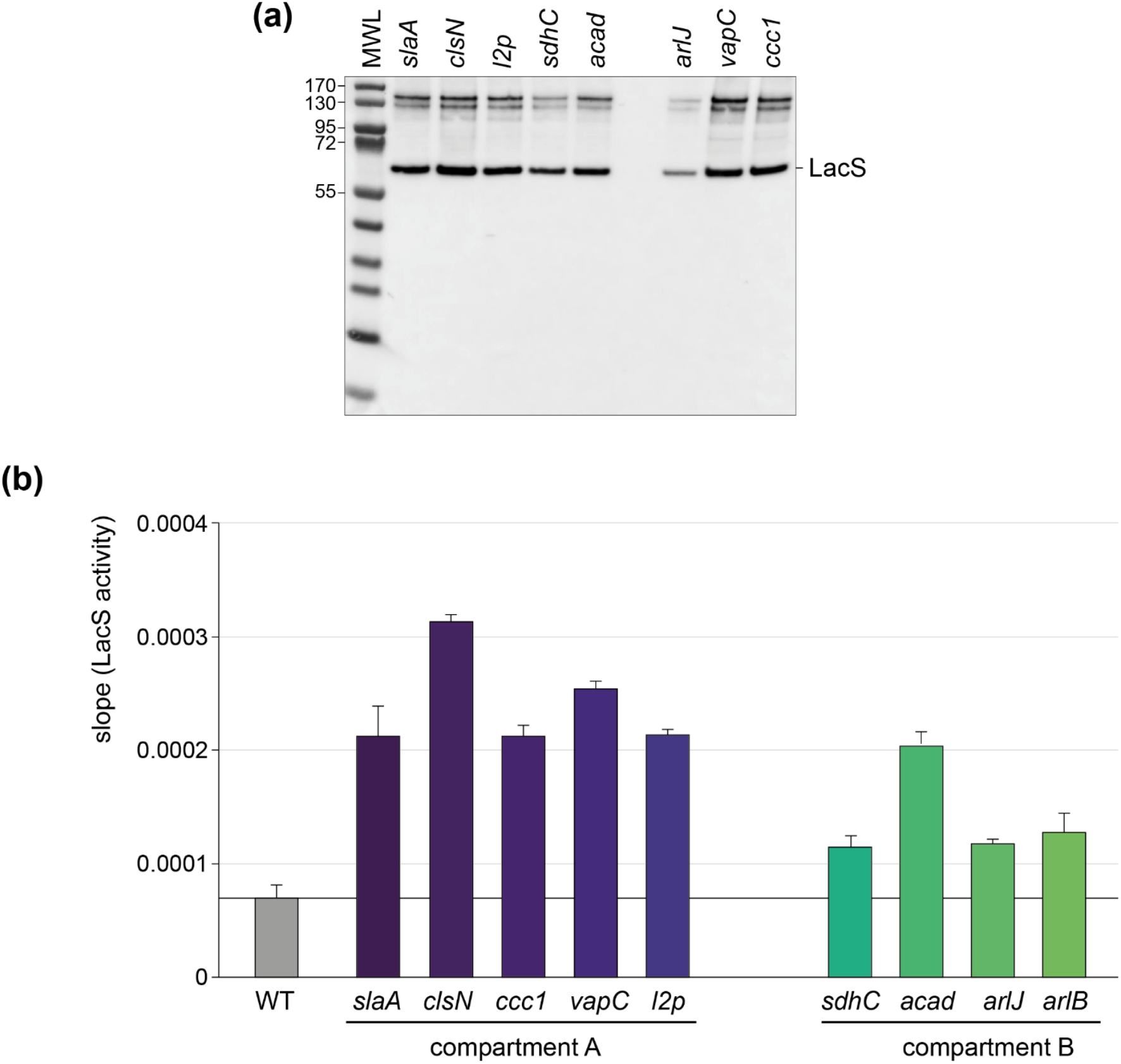
Translational and activity levels of LacS. (**a**) Western blot analysis employing anti-6xHis antibodies for the detection of LacS-6xHis in the different knock-in strains. The *arlB* knock-in strain was not included in this analysis because of a mutation in the His-tag-encoding sequence. MWL = molecular weight ladder. Protein molecular weights are indicated in kDa. (**b**) LacS activities as measured with an ONPG assay for the different knock-in strains.

The observed differences in translational activity for the different strains were less pronounced as those observed at the transcriptional level. For instance, while compartment A strains, such as the *clsN* and *slaA* knock-in strains, exhibited slightly higher protein activities as compared to compartment B strains, such as the *arlJ* knock-in strain, the variation in protein activity was relatively modest. Moreover, the correlation between transcriptional and translational levels was limited with a *R*^2^ of 0.2358 (**Figure 7a**). Interestingly, a larger correlation can be found for protein activity and distance to the closest origin of replication (*R*^2^ = 0.432) than for transcriptional expression (**Figures 7b** and **7c**).

**Figure 7.**
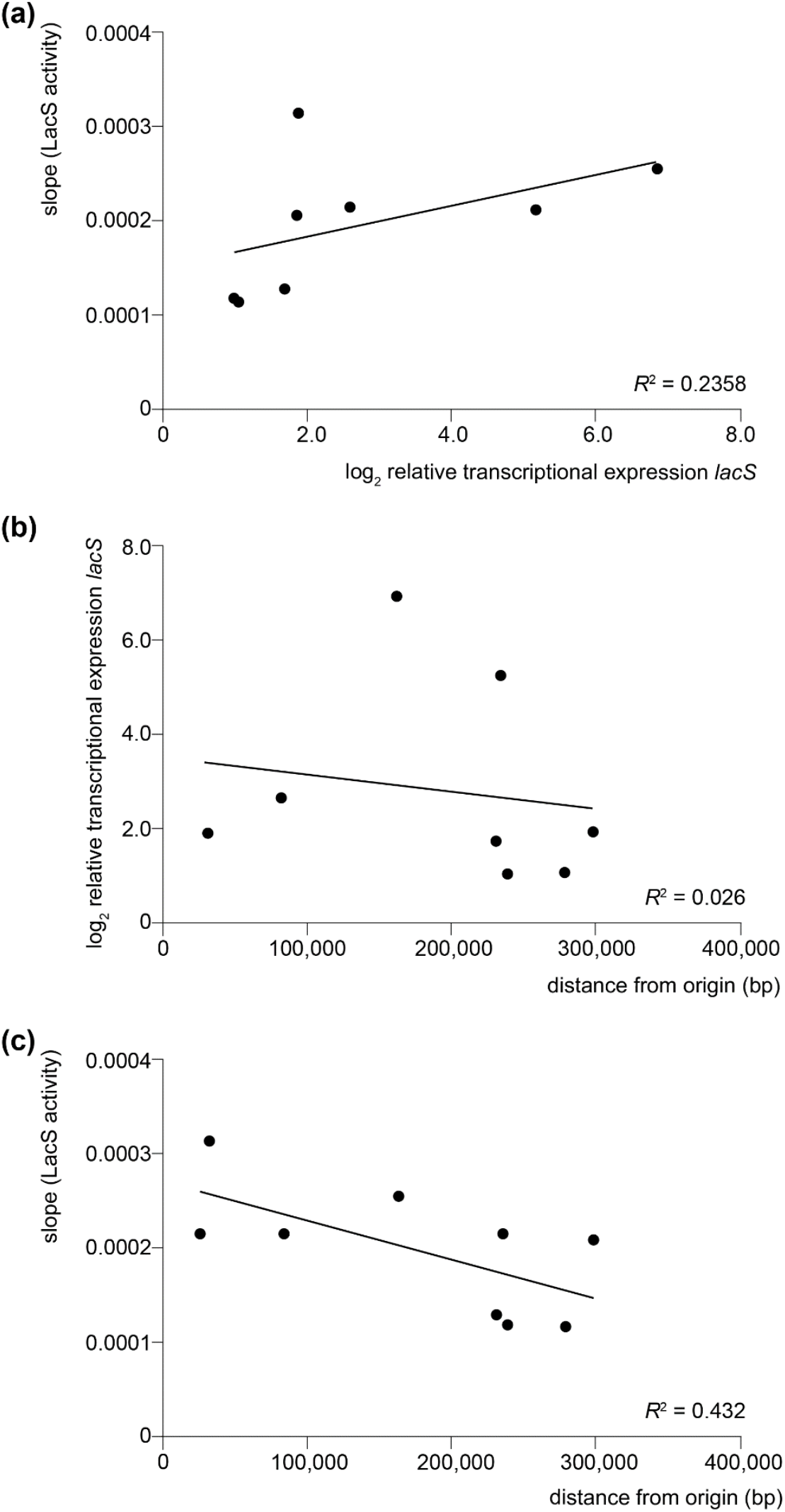
Correlation between transcription, translation and distance to origin of replication. (**a**) Scatter plot between translational levels as monitored by LacS activities and relative transcriptional expression levels of *lacS* as monitored by qRT-PCR analysis. The data point corresponding to the *slaA* knock-in strain was omitted given the exceptionally high relative transcriptional level. (**b**) Scatter plot between relative transcriptional expression levels of *lacS* as monitored by qRT-PCR analysis and distance to the closest origin of replication. The data point corresponding to the *slaA* knock-in strain was omitted given the exceptionally high relative transcriptional level. (**c**) Scatter plot between translational levels as monitored by LacS activities and distance to the closest origin of replication. For all plots, a linear regression line has been fitted and the coefficient of determination (*R*^2^) is displayed.

## 4 Discussion

In conclusion, our study indicates that while genomic location can influence gene expression levels in *S. acidocaldarius*, the evidence for a strong position-dependent effect independent of local genomic context is limited, especially at the transcriptional level. Instead, transcriptional variability can be largely attributed to the transcriptional activity of neighboring genes rather than to the genomic position itself for most of the chosen target positions. Indeed, these positions did not insulate the *lacS* cassette from surrounding transcriptional influences, leading to expression levels that were primarily dictated by local transcriptional context. This can be explained by the high coding density in the *S. acidocaldarius* genome, with 2,292 protein-encoding genes predicted for 2,225,959 bp (Chen et al. 2005). Therefore, while almost all target sites are in intergenic regions, only in the *clsN* and *acad* knock-in strains these target sites can be assumed not to be part of a transcriptional unit. Moreover, due to a lack of the availability of genetic parts for archaea, for example a transcriptional terminator, we could not integrate an insulator element into the reporter gene cassette as is typically done in bacteria (Scholz et al. 2022).

The strong influence of the activity of adjacent transcription units is not only apparent for constitutive expression levels, but also for transcription regulation. For most knock-in strains, similar regulatory trends were observed for two different stress conditions: nutritional starvation and stationary phase growth. Moreover, the regulatory effect typically corresponds to that of the adjacent gene. For example, this is the case for the *arlB* knock-in strain. The *arlB* gene encodes the main structural unit of the archaellum and is part of a larger operon that encodes additional structural and functional proteins of this motility structure (Lassak et al, 2012). This operon is known to be transcriptionally upregulated in response to stress conditions such as nutrient limitation, through a complex interplay of different transcription regulators and kinases (Lassak et al, 2013, Haurat et al, 2017, Li et al, 2017). By incorporating the reporter gene cassette directly downstream of the *arlB* ORF, it is also subject to this regulation. This was most clearly observed in response to nutritional starvation, but also in response to stationary phase growth, albeit less pronounced.

There was a lack of a clear correlation between transcriptional and translational levels. This suggests that translational efficiency and LacS activity may be influenced by factors beyond transcription alone, such as mRNA stability and ribosome accessibility. This lack of correlation between transcription and translation was previously observed for dynamic effects in response to nutrient limitation (Bischof et al. 2018) and heat shock (Baes et al. 2023), indicating a prevalence of post-transcriptional and post-translational mechanisms in *S. acidocaldarius*. Interestingly, translational activity exhibited a more apparent correlation with the distance to the closest origin of replication as compared to transcriptional activity.

Several of the ectopic integration sites explored in our study could be employed in the future for the construction of knock-in strains, whether it is for the generation of strains with altered phenotypic characteristics or for high-level production of recombinant protein. It should be noted that in our study, the *lacS* ORF was directly fused to a promoter, without including a 5’-untranslated region (5’-UTR). Although the presence of a Shine-Dalgarno (SD)-containing 5’-UTR sequence might be considered less relevant for gene expression, given the prevalent occurrence of leaderless transcripts in archaea (Wurtzel et al, 2010, Schmitt et al, 2020), it has been shown that the presence of a 5’-UTR in plasmid-based heterologous expression constructs in Sulfolobales can significantly impact the gene expression level by affecting both transcriptional and translational efficiency (Peng et al, 2009, Ao et al, 2013, Kuschmierz et al, 2024). More specifically, the integration of a 5’-UTR derived from the Alba-encoding gene *saci_1322* from *S. acidocaldarius* harboring a SD sequence in a heterologous protein expression plasmid led to a significantly higher protein production (Kuschmierz et al, 2024). Therefore, if the goal is to obtain a maximal yield of recombinant protein production in a stable *S. acidocaldarius* platform, it could be advised to integrate the *alba*-derived 5’-UTR into an expression cassette ectopically integrated into the genome target site just downstream of the *slaA* ORF.

## Conflict of Interest

The authors declare that the research was conducted in the absence of any commercial or financial relationships that could be construed as a potential conflict of interest.

## Author Contributions

Y.X.: Conceptualization, Investigation, Formal analysis, Writing – original draft. A.I.P.: Investigation, Formal analysis, Writing – review & editing. I.B.: Supervision, Writing – review & editing. M.D.M.: Resources, Supervision, Writing – review & editing. R.B.: Formal analysis, Supervision, Writing – review & editing. E.P.: Conceptualization, Resources, Supervision, Visualization, Writing – original draft.

## Funding

This research was funded by the Vrije Universiteit Brussel (Strategic Research Program SRP91), by Flanders Innovation and Entrepreneurship (VLAIO) (Moonshot project “TACBIO” [HBC.2020.2618]), by the Bijzonder Onderzoeksfonds (iBOF project “POSSIBL” [iBOF/21/092]) and by the Research Foundation Flanders (FWO-Vlaanderen) [G012323N]. Y.X. was funded by a PhD scholarship from the Chinese Scholarship Council.

## Acknowledgments

We would like to thank Norio Kurosawa for the generous gift of the *S. acidocaldarius* SK-1 strain and Karl Jonckheere for technical assistance.

## Generative AI statement

The authors declare that no Gen AI was used in the creation of this manuscript.

## Data Availability Statement

Data are available on request.

## Notes

### Competing Interest Statement

The authors have declared no competing interest.

