## Supplementary for "Effects of genomic location on ectopic integration and gene expression of a reporter gene cassette in *Sulfolobus acidocaldarius*"

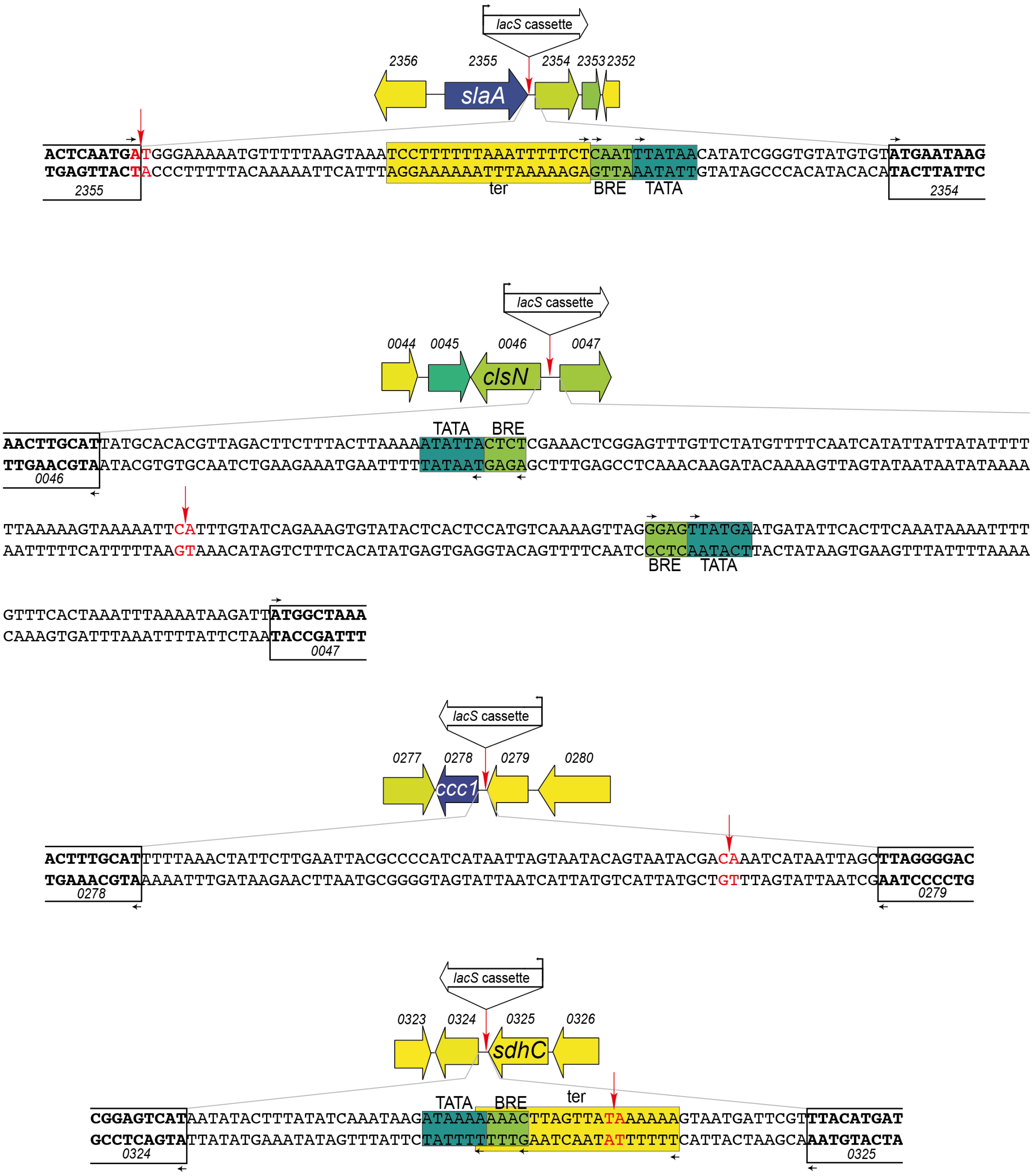


**Supplementary Figure S1.** Sequences of the genomic environment for each of the target sites for integration of the *lacS* cassette. The schematic depiction displays the target site and the orientation of the *lacS* cassette with respect to the genomic environment. Numbers refer to gene locus tags, with xxxx referring to *Saci_xxxx*. Gene arrows are color-coded according to expression levels (CPM value) (Baes et al. 2023) according to the color scheme shown in **Figure 3b**. On the sequences, the target site is indicated in red, ORFs are boxed with the direction of transcription indicated by an arrow. Putative promoter elements are indicated with TATA = TATA box and BRE = factor B recognition element and putative terminator elements are indicated by ter. Arrows indicate the direction of transcription of the corresponding transcription unit.

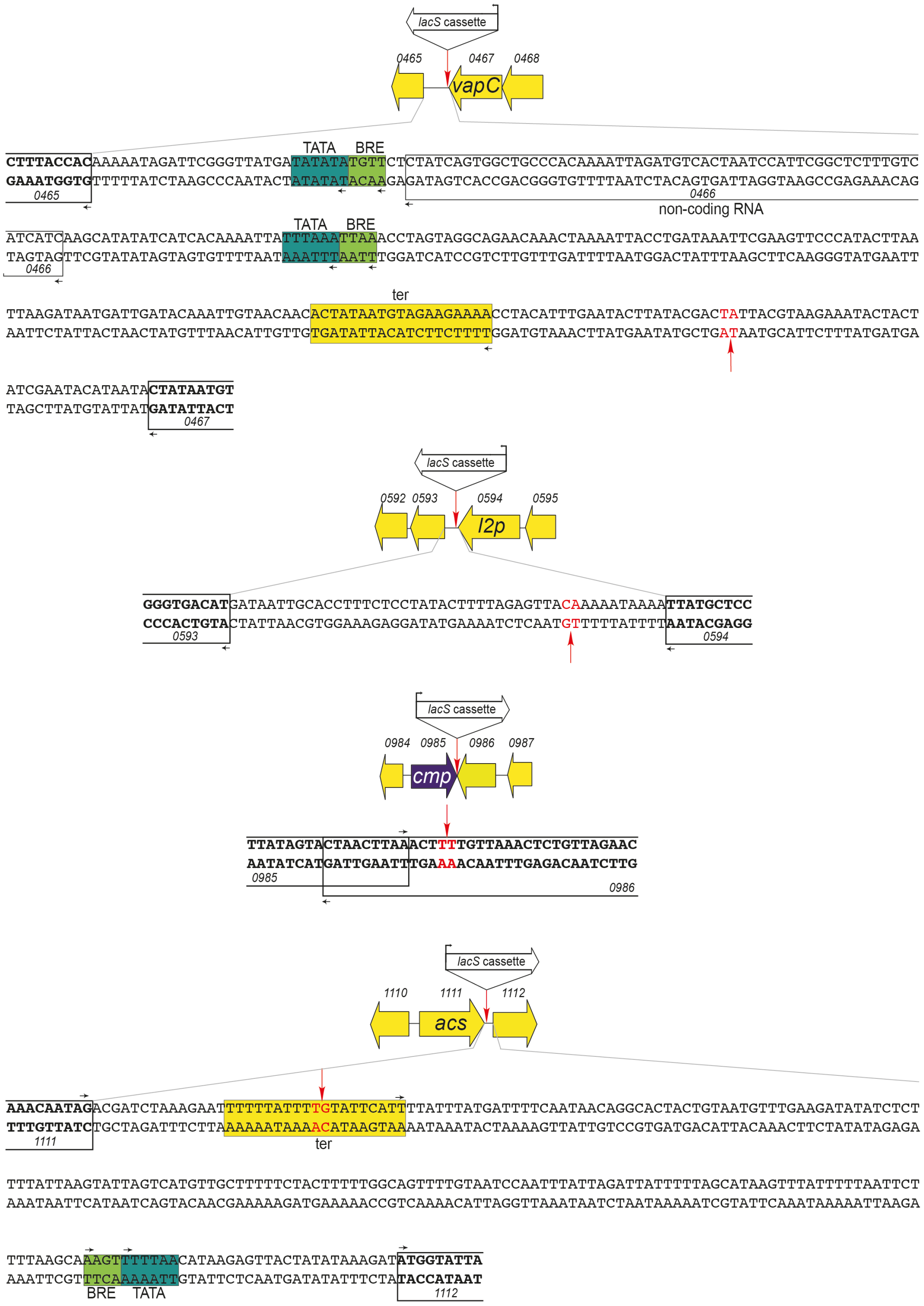

**Supplementary Figure S1.** *Continued.*

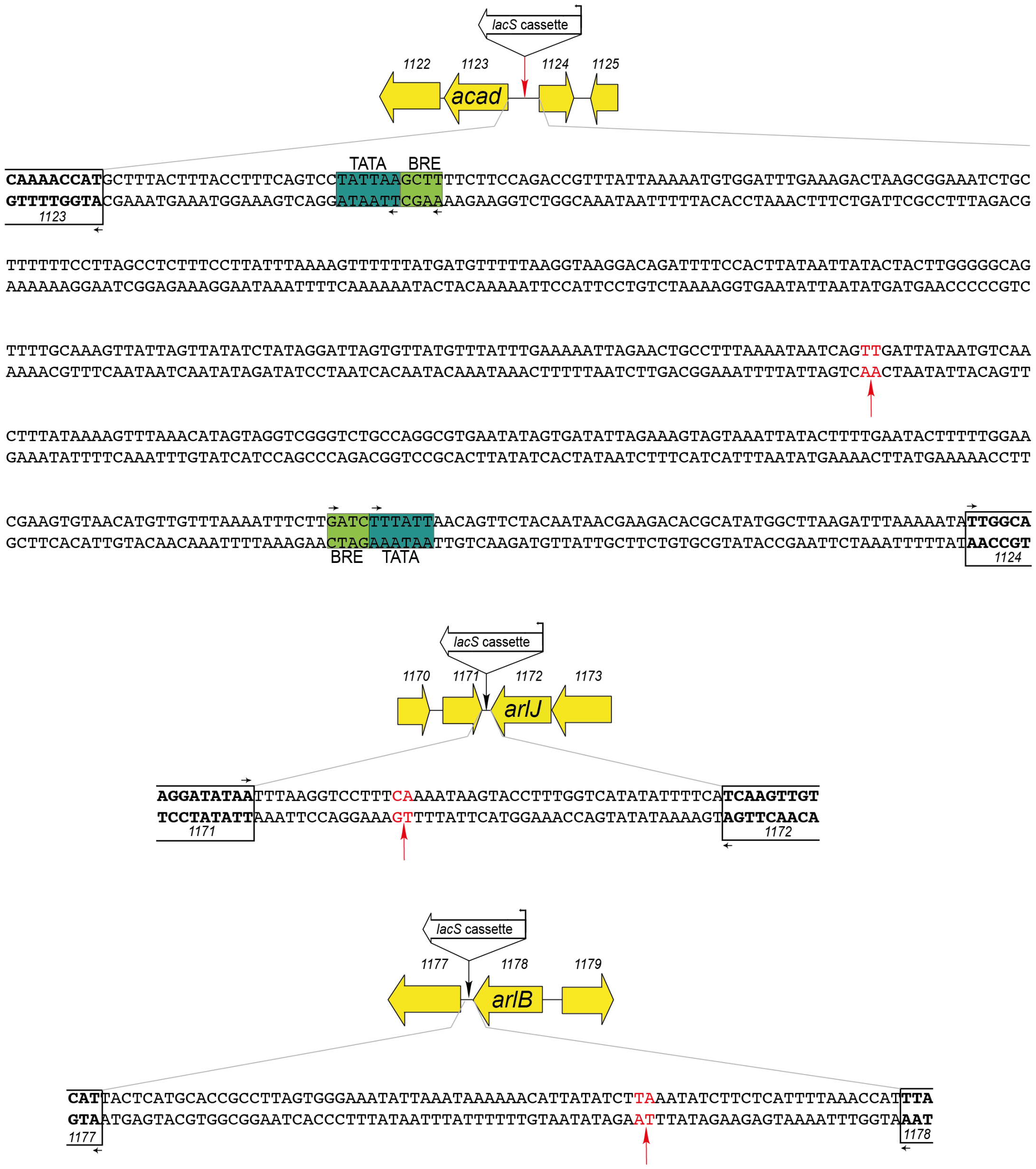


**Supplementary Figure S1.** *Continued.*

**Supplementary Table S1.** Overview of all microbial strains used in this work.

| Name | Description | Origin |
|---|---|---|
| *Escherichia coli* DH5α | Host strain for cloning and plasmid propagation | Life Technologies |
| *Sulfolobus acidocaldarius* SK-1 | Uracil-auxotrophic host strain | (Suzuki and Kurosawa 2016) |
| *S. acidocaldarius* SK1-*slaAx*KI*lacs* | Knock-in strain with a *lacS* cassette in the *slaA* locus | This study |
| *S. acidocaldarius* SK1-*clsNx*KI*lacs* | Knock-in strain with a *lacS* cassette in the *clsN* locus | This study |
| *S. acidocaldarius* SK1-*l2px*KI*lacs* | Knock-in strain with a *lacS* cassette in the *l2p* locus | This study |
| *S. acidocaldarius* SK1-*sdhCx*KI*lacs* | Knock-in strain with a *lacS* cassette in the *sdhC* locus | This study |
| *S. acidocaldarius* SK1-*fad11x*KI*lacs* | Knock-in strain with a *lacS* cassette in the *fad11* locus | This study |
| *S. acidocaldarius* SK1-*fad23x*KI*lacs* | Knock-in strain with a *lacS* cassette in the *fad23* locus | This study |
| *S. acidocaldarius* SK1-*flaBx*KI*lacs* | Knock-in strain with a *lacS* cassette in the *flaB* locus | This study |
| *S. acidocaldarius* SK1-*flaJx*KI*lacs* | Knock-in strain with a *lacS* cassette in the *flaJ* locus | This study |
| *S. acidocaldarius* SK1-*vapCx*KI*lacs* | Knock-in strain with a *lacS* cassette in the *vapC* locus | This study |
| *S. acidocaldarius* SK1-*cmpx*KI*lacs* | Knock-in strain with a *lacS* cassette in the *cmp* locus | This study |
| *S. acidocaldarius* SK1-*ccc1x*KI*lacs* | Knock-in strain with a *lacS* cassette in the *ccc1* locus | This study |

**Supplementary Table S2**. Overview of oligonucleotides used in this work, with indication of their sequence and purpose. Golden Gate linker sequences are depicted in bold, with linker 1 = cut site 5'-AGGA-3', linker 2 = cut site 5'-TGAT-3', linker 3 = cut site 5'-GATG-3' and linker 4 = cut site 5'-GCAG-3'. For primer YX091, the sequence of the SP1 promoter is indicated in bold.

| Name | Sequence (5' – 3') | Purpose |
|---|---|---|
| YX091 | **TATAATACAAATCACTCTCAAATGGTTTTATAAACTGAGGGGAGAGAATATTACTT**ATGGACTCATTTCCAAATAGCTTTAGGTTTG | Switch in pSVA431 from P*malE* to SP1 promoter. |
| YX123 | CACCTGCCGATGCAGAATTCGCCCTATAGTGAGTCGTATTACAAT | Used with YX124 to amplify the backbone of pSVA431. |
| YX124 | CACCTGCCGATTCCTGATAAGCATGCATGACCGGCTATT | Used with YX123 to amplify the backbone of pSVA431. |
| YX125 | TCATGCATGCTTATCAGGAATCGGCAGGTGC | Used with YX126 to amplify the *sacB* gene. |
| YX126 | ACTATAGGGCGAATTCTGCATCGGCAGGTGATGGGAT | Used with YX127 to amplify the *sacB* gene. |
| YX127 | ATAAGGTGATGAAATGTAAAGGAGC | Used with YX128 for performing colony PCR and sequencing inserts in pYX2304. |
| YX128 | GTAAAACGACGGCCAGT | Used with YX127 for performing colony PCR and sequencing inserts in pYX2304. |
| YX131 | TATTCACCTGCACTA**AGGA**AGACTTCATTTGCAGTTTACACCAAT | Used with YX132 to amplify the upstream region of *slaA* locus, to assemble pYX2304-1. It contains a linker 1 for Golden Gate cloning. |
| YX132 | TATTCACCTGCACTA**ATCA**TTGAGTTAATGTTACGTTTGTGAATATTAATTGT | Used with YX131 to amplify the upstream region of *slaA* locus, to assemble pYX2304-1. It contains a linker 2 for Golden Gate cloning. |
| YX133 | TATTCACCTGCACTA**TGAT**ATAATACAAATCACTCTCAAATGGTTTTATAAACTGAGGG | Used with YX134 to amplify the *lacS* cassette, to assemble all pYX2304-based plasmids. It contains a linker 2 for Golden Gate cloning. |
| YX134 | TATTCACCTGCACTA**CATC**AGTGGTGGTGGTGGTGGTGGTGCCTTAATGGCTTTACTGGA | Used with YX133 to amplify the *lacS* cassette, to assemble all pYX2304-based plasmids. It contains a linker 3 for Golden Gate cloning. |
| YX135 | TATTCACCTGCACTA**GATG**GGAAAAAATGTTTTTAAGTAAATCCTTTTTTAAATTTTTCTC | Used with YX136 to amplify the downstream region of *slaA* locus, to assemble pYX2304-1. It contains a linker 3 for Golden Gate cloning. |
| YX136 | TATTCACCTGCACTA**CTGC**CTTGAAGCGAAGCTACTAACTTAAATGT | Used with YX135 to amplify the downstream region of *slaA* locus, to assemble pYX2304-1. It contains a linker 4 for Golden Gate cloning. |
| YX137 | AATTAGTGAAAAAATAGCCGGTC | Used with YX138 for sequencing of all pYX2304-based plasmid constructs. |
| YX138 | GTAAAACGACGGCCAGT | Used with YX137 for sequencing of all pYX2304-based plasmid constructs. |
| YX139 | TATAATACAAATCACTCTCAAATGG | Used with YX140 for sequencing of all pYX2304-based plasmid constructs. |
| YX140 | GTGCCTTAATGGCTTTACTG | Used with YX139 for sequencing of all pYX2304-based plasmid constructs. |
| YX141 | TATTCACCTGCACTA**AGGA**CTTTTCCTTTTCGAGCCTTTCAATTTCCA | Used with YX142 to amplify the upstream region of *clsN* locus, to assemble pYX2304-2. It contains a linker 1 for Golden Gate cloning. |

| | | |
|---|---|---|
| YX142 | TATTCACCTGCACTA**ATCA**GAATTTTTACTTTTTAAAAAATATAATAATATGATTGAAAACATAGAACA | Used with YX141 to amplify the upstream region of *clsN* locus, to assemble pYX2304-2. It contains a linker 2 for Golden Gate cloning. |
| YX143 | TATTCACCTGCACTA**GATG**ATTTGTATCAGAAAGTGTATACTCACTCCATGT | Used with YX144 to amplify the downstream region of *clsN* locus, to assemble pYX2304-2. It contains a linker 3 for Golden Gate cloning. |
| YX144 | TATTCACCTGCACTA**CTGC**ATCAAACCCAGCTGAAACTACGATATTT | Used with YX143 to amplify the downstream region of *clsN* locus, to assemble pYX2304-2. It contains a linker 4 for Golden Gate cloning. |
| YX145 | TATTCACCTGCACTA**AGGA**GCCAGCTAGATATCCAAATATAGAGGGAGA | Used with YX146 to amplify the upstream region of *l2p* locus, to assemble pYX2304-3. It contains a linker 1 for Golden Gate cloning. |
| YX146 | TATTCACCTGCACTA**ATCA**AAAAATAAAATTATGCTCCACCACGTCTACC | Used with YX145 to amplify the upstream region of *l2p* locus, to assemble pYX2304-3. It contains a linker 2 for Golden Gate cloning. |
| YX147 | TATTCACCTGCACTA**GATG**GTAACTCTAAAAGTATAGGAGAAAGGTGCAATTATCA | Used with YX148 to amplify the downstream region of *l2p* locus, to assemble pYX2304-3. It contains a linker 3 for Golden Gate cloning. |
| YX148 | TATTCACCTGCACTA**CTGC**CAGAACGGTAAAGCTTCCTT | Used with YX147 to amplify the downstream region of *l2p* locus, to assemble pYX2304-3. It contains a linker 4 for Golden Gate cloning. |
| YX149 | TATTCACCTGCACTA**AGGA**AATACTACATGTCAAAATATCCTAAGGTCAGAGAGA | Used with YX150 to amplify the upstream region of *sdhC* locus, to assemble pYX2304-4. It contains a linker 1 for Golden Gate cloning. |
| YX150 | TATTCACCTGCACTA**ATCA**AAAAAAGTAATGATTCGTTTACATGTATCTCTTTAACACTG | Used with YX149 to amplify the upstream region of *sdhC* locus, to assemble pYX2304-4. It contains a linker 2 for Golden Gate cloning. |
| YX151 | TATTCACCTGCACTA**GATG**ATAACTTAAGTTTTTTTATCTTATTTGATATAAAGTATATTATGACTCCG | Used with YX152 to amplify the downstream region of *sdhC* locus, to assemble pYX2304-4. It contains a linker 3 for Golden Gate cloning. |
| YX152 | TATTCACCTGCACTA**CTGC**AATGATACTCATAGAGGTACTAAAGAGTAATGCAGGAAC | Used with YX151 to amplify the downstream region of *sdhC* locus, to assemble pYX2304-4. It contains a linker 4 for Golden Gate cloning. |
| YX153 | TATTCACCTGCACTA**AGGA**CTAAGCCCGAATTCGAGCA | Used with YX154 to amplify the upstream region of *fad11* locus, to assemble pYX2304-5. It contains a linker 1 for Golden Gate cloning. |
| YX154 | TATTCACCTGCACTA**ATCA**AAAATAAAAAATTCTTTAGATCGTCTATTGTTTCATG | Used with YX153 to amplify the upstream region of *fad11* locus, to assemble pYX2304-5. It contains a linker 2 for Golden Gate cloning. |
| YX155 | TATTCACCTGCACTA**GATG**GTATTCATTTTATTTATGATTTTCAATAACAGGCAC | Used with YX156 to amplify the downstream region of *fad11* locus, to assemble pYX2304-5. It contains a linker 3 for Golden Gate cloning. |
| YX156 | TATTCACCTGCACTA**CTGC**TGCCTTTGGGAGCTCCT | Used with YX155 to amplify the downstream region of *fad11* locus, to assemble pYX2304-5. It contains a linker 4 for Golden Gate cloning. |
| YX157 | TATTCACCTGCACTA**AGGA**GATAGTCAGTGAACCTTTCCG | Used with YX158 to amplify the upstream region of *fad23* locus, to assemble pYX2304-6. It contains a linker 1 for Golden Gate cloning. |
| YX158 | TATTCACCTGCACTA**ATCA**TGATTATAATGTCAACTTTATAAAAGTTTAAACATAGTAGG | Used with YX157 to amplify the upstream region of *fad23* locus, to assemble pYX2304-6. It contains a linker 2 for Golden Gate cloning. |

| | | |
|---|---|---|
| YX159 | TATTCACCTGCACTA**GATG**ACTGATTATT TTAAAGGCAGTTCTAATTTTTCA | Used with YX160 to amplify the downstream region of *fad23* locus, to assemble pYX2304-6. It contains a linker 3 for Golden Gate cloning. |
| YX160 | TATTCACCTGCACTA**CTGC**CTTATCTCCC TTGGCAACAG | Used with YX159 to amplify the downstream region of *fad23* locus, to assemble pYX2304-6. It contains a linker 4 for Golden Gate cloning. |
| YX161 | TATTCACCTGCACTA**AGGA**TATCTCCTTC AACTACAGCAATATCG | Used with YX162 to amplify the upstream region of *flaB* locus, to assemble pYX2304-7. It contains a linker 1 for Golden Gate cloning. |
| YX162 | TATTCACCTGCACTA**ATCA**AAATATCTTC TCATTTTAAACCATTTATCCTATAACTG | Used with YX161 to amplify the upstream region of *flaB* locus, to assemble pYX2304-7. It contains a linker 2 for Golden Gate cloning. |
| YX163 | TATTCACCTGCACTA**GATG**AAGATATAAT GTTTTTTATTTAATATTTCCCACTAAGGC | Used with YX164 to amplify the downstream region of *flaB* locus, to assemble pYX2304-7. It contains a linker 3 for Golden Gate cloning. |
| YX164 | TATTCACCTGCACTA**CTGC**GTAAACATAT CATCTGAAGAGACATATCCTAATTC | Used with YX163 to amplify the downstream region of *flaB* locus, to assemble pYX2304-7. It contains a linker 4 for Golden Gate cloning. |
| YX165 | TATTCACCTGCACTA**AGGA**TAAGGGGTGA TTTAGGTTCTGC | Used with YX166 to amplify the upstream region of *flaJ* locus, to assemble pYX2304-8. It contains a linker 1 for Golden Gate cloning. |
| YX166 | TATTCACCTGCACTA**ATCA**AAAATAAGTA CCTTTGGTCATATATTTTCATCAAG | Used with YX165 to amplify the upstream region of *flaJ* locus, to assemble pYX2304-8. It contains a linker 2 for Golden Gate cloning. |
| YX167 | TATTCACCTGCACTA**GATG**GAAAGGACCT TAAATTATATCCTGTTAAATATTTCT | Used with YX168 to amplify the downstream region of *flaJ* locus, to assemble pYX2304-8. It contains a linker 3 for Golden Gate cloning. |
| YX168 | TATTCACCTGCACTA**CTGC**GCAATAAGAA AACTAGTTGACAAAGAAC | Used with YX167 to amplify the downstream region of *flaJ* locus, to assemble pYX2304-8. It contains a linker 4 for Golden Gate cloning. |
| YX169 | TATTCACCTGCACTA**AGGA**TAAATATATT TATGCCTAAAGTCATTACAATATCAGATG | Used with YX170 to amplify the upstream region of *vapC* locus, to assemble pYX2304-9. It contains a linker 1 for Golden Gate cloning. |
| YX170 | TATTCACCTGCACTA**ATCA**ATTACGTAAG AAATACTACTATCGAATACATAATACTAT AAT | Used with YX169 to amplify the upstream region of *vapC* locus, to assemble pYX2304-9. It contains a linker 2 for Golden Gate cloning. |
| YX171 | TATTCACCTGCACTA**GATG**AGTCGTATAA GTATTCAAATGTAGGTTTTCT | Used with YX172 to amplify the downstream region of *vapC* locus, to assemble pYX2304-9. It contains a linker 3 for Golden Gate cloning. |
| YX172 | TATTCACCTGCACTA**CTGC**TTAGTATCCT CGAGTAGTTAAGAATCGAC | Used with YX171 to amplify the downstream region of *vapC* locus, to assemble pYX2304-9. It contains a linker 4 for Golden Gate cloning. |
| YX173 | TATTCACCTGCACTA**AGGA**GTTATGGGAG AACTCAGTAAAGTATAATG | Used with YX174 to amplify the upstream region of *cmp* locus, to assemble pYX2304-10. It contains a linker 1 for Golden Gate cloning. |
| YX174 | TATTCACCTGCACTA**ATCA**AAGTTTAAGT TAGTACTATAAAAATAATTGCAAGAAATA G | Used with YX173 to amplify the upstream region of *cmp* locus, to assemble pYX2304-10. It contains a linker 2 for Golden Gate cloning. |
| YX175 | TATTCACCTGCACTA**GATG**TTGTTAAACT CTGTTAGAACATTGTCAACTT | Used with YX176 to amplify the downstream region of *cmp* locus, to assemble pYX2304-10. It contains a linker 3 for Golden Gate cloning. |

| | | |
|---|---|---|
| YX176 | TATTCACCTGCACTA**CTGC**CAGGGCGGTATGGGAT | Used with YX175 to amplify the downstream region of *cmp* locus, to assemble pYX2304-10. It contains a linker 4 for Golden Gate cloning. |
| YX177 | TATTCACCTGCACTA**AGGA**TACATTTAACTTGGAGTTGTTACGAGG | Used with YX178 to amplify the upstream region of *ccc1* locus, to assemble pYX2304-11. It contains a linker 1 for Golden Gate cloning. |
| YX178 | TATTCACCTGCACTA**ATCA**AAATCATAATTAGCTTAGGGGACAGC | Used with YX177 to amplify the upstream region of *ccc1* locus, to assemble pYX2304-11. It contains a linker 2 for Golden Gate cloning. |
| YX179 | TATTCACCTGCACTA**GATG**GTCGTATTACTGTATTACTAATTATGATGGG | Used with YX180 to amplify the downstream region of *ccc1* locus, to assemble pYX2304-11. It contains a linker 3 for Golden Gate cloning. |
| YX180 | TATTCACCTGCACTA**CTGC**TAACAAGTATAACAGCCATTGAGGAC | Used with YX179 to amplify the downstream region of *ccc1* locus, to assemble pYX2304-11. It contains a linker 4 for Golden Gate cloning. |
| YX183 | TTAAGCGCAGTACCATTTGC | Used with YX184 for colony PCR of *S. acidocaldarius* SK1-*slaAx*KI*lacs*. |
| YX184 | GCTAAGTTCTTATTCATACACATACAC | Used with YX183 for colony PCR of *S. acidocaldarius* SK1-*slaAx*KI*lacs*. |
| YX185 | TCTAGCGATAATACTAGCTCTC | Used with YX186 for colony PCR of *S. acidocaldarius* SK1-*clsNx*KI*lacs*. |
| YX186 | GACATGGAGTGAGTATACAC | Used with YX185 for colony PCR of *S. acidocaldarius* SK1-*clsNx*KI*lacs*. |
| YX187 | GTCATATTGCATCAAGAAGGAC | Used with YX188 for colony PCR of *S. acidocaldarius* SK1-*l2px*KI*lacs*. |
| YX188 | GATAATTGCACCTTTCTCCTA | Used with YX187 for colony PCR of *S. acidocaldarius* SK1-*l2px*KI*lacs*. |
| YX189 | CCTATGTCAGTGTTAAAGAGATAC | Used with YX190 for colony PCR of *S. acidocaldarius* SK1-*sdhCx*KI*lacs*. |
| YX190 | GTTAATTCCACCTCAGTATACG | Used with YX189 for colony PCR of *S. acidocaldarius* SK1-*sdhCx*KI*lacs*. |
| YX191 | GGAGTTGAGAAATAAGTATAAGGAC | Used with YX192 for colony PCR of *S. acidocaldarius* SK1-*fad11x*KI*lacs*. |
| YX192 | GTAGTGCCTGTTATTGAAAATC | Used with YX191 for colony PCR of *S. acidocaldarius* SK1-*fad11x*KI*lacs*. |
| YX193 | ACAACATGTTACACTTCGTATC | Used with YX194 for colony PCR of *S. acidocaldarius* SK1-*fad23x*KI*lacs*. |
| YX194 | CGGAAATCTGCTTTTTTTCC | Used with YX193 for colony PCR of *S. acidocaldarius* SK1-*fad23x*KI*lacs*. |
| YX195 | CAATATCCCAGTATATCTATCAGC | Used with YX196 for colony PCR of *S. acidocaldarius* SK1-*flaBx*KI*lacs*. |
| YX196 | CTTGGATAGCCATTACTCAT | Used with YX195 for colony PCR of *S. acidocaldarius* SK1-*flaBx*KI*lacs*. |
| YX197 | CTGGAGGTATAATTAGTGGC | Used with YX198 for colony PCR of *S. acidocaldarius* SK1-*flaJx*KI*lacs*. |
| YX198 | CGTTACTTTCGATATTCGCA | Used with YX197 for colony PCR of *S. acidocaldarius* SK1-*flaJx*KI*lacs*. |
| YX199 | AAGTCTAATAATGAGGTTTTGATAAC | Used with YX200 for colony PCR of *S. acidocaldarius* SK1-*vapCx*KI*lacs*. |
| YX200 | GATTGATACAAATTGTAACAACACT | Used with YX199 for colony PCR of *S. acidocaldarius* SK1-*vapCx*KI*lacs*. |

| | | |
|---|---|---|
| YX201 | GGAGGTTATGTTATGAAAACATTC | Used with YX202 for colony PCR of *S. acidocaldarius* SK1-*cmpx*KI*lacs*. |
| YX202 | CTTTAGGTAGTTGGAGAAGAGA | Used with YX201 for colony PCR of *S. acidocaldarius* SK1-*cmpx*KI*lacs*. |
| YX203 | AATGGGGTATATACTCCAATTTC | Used with YX204 for colony PCR of *S. acidocaldarius* SK1-*cc1x*KI*lacs*. |
| YX204 | GAGTCCATCTTGTATTCCG | Used with YX203 for colony PCR of *S. acidocaldarius* SK1-*cc1x*KI*lacs*. |

**Supplementary Table S3.** Overview of all plasmids used in this work.

| Name | Description | Origin |
|---|---|---|
| pSVA431 | Suicide vector for construction of mutant strains | (Wagner et al. 2012) |
| pYX2301-UPDD | pSVA431 variant with SP1 promoter | This study |
| pRN1-carrier_2_paqci | Construction of Golden Gate destination vector | This study |
| pYX2304 | Golden Gate destination vector | This study |
| pYX2304-1 | Suicide vector for construction of *S. acidocaldarius* SK1-*slaAx*KI*lacs* | This study |
| pYX2304-2 | Suicide vector for construction of *S. acidocaldarius* SK1-*clsNx*KI*lacs* | This study |
| pYX2304-3 | Suicide vector for construction of *S. acidocaldarius* SK1-*l2px*KI*lacs* | This study |
| pYX2304-4 | Suicide vector for construction of *S. acidocaldarius* SK1-*sdhCx*KI*lacs* | This study |
| pYX2304-5 | Suicide vector for construction of *S. acidocaldarius* SK1-*fad11Ax*KI*lacs* | This study |
| pYX2304-6 | Suicide vector for construction of *S. acidocaldarius* SK1-*fad23x*KI*lacs* | This study |
| pYX2304-7 | Suicide vector for construction of *S. acidocaldarius* SK1-*flaBx*KI*lacs* | This study |
| pYX2304-8 | Suicide vector for construction of *S. acidocaldarius* SK1-*flaJx*KI*lacs* | This study |
| pYX2304-9 | Suicide vector for construction of *S. acidocaldarius* SK1-*vapCx*KI*lacs* | This study |
| pYX2304-10 | Suicide vector for construction of *S. acidocaldarius* SK1-*cmpx*KI*lacs* | This study |
| pYX2304-11 | Suicide vector for construction of *S. acidocaldarius* SK1-*ccc1x*KI*lacs* | This study |

**Supplementary Table S4.** Comparison between relative transcriptional expression level of *lacS* as measured by qRT-PCR in the different knock-in strains, relative to the *arlJ* knock-in strain (**Figure 5a**) and expression levels of the adjacent gene of which the transcription unit is most closely located to the *lacS* cassette, expressed in CPM value (Baes et al. 2023).

| **Knock-in strain** | **$Log_2$ fold ratio *lacS* expression** | **Adjacent gene of which the transcription unit is most closely located to the *lacS* cassette** | **Expression level** |
|---|---|---|---|
| *slaA* | 5.77 | *saci_2355* | 11554 |
| *ccc1* | 2.29 | *saci_0279* | 55012 |
| *sdhC* | 0.00 | *saci_0325* | 129.5 |
| *vapC* | 2.28 | *saci_0467* | 1.47 |
| *l2p* | 1.16 | *saci_0594* | 53.09 |
| *arlJ* | 0.00 | *saci_1172* | 257.89 |
| *arlB* | 0.62 | *saci_1178* | 35.18 |
